## Supplementary Figures 1-13 for "An Ensemble Penalized Regression Method for Multi-ancestry Polygenic Risk Prediction"

**Supplementary Figure 1: Optimal tuning parameter lambda in lasso.** The simulation is performed for design matrix with $p = 1000$ predictors, and 5% of them are randomly selected to be causal. Correlation structure of those predictors is AR1 with $\rho = 0.4$. The total heritability is simulated to be $0.2$. Source data are provided as a Source Data file.

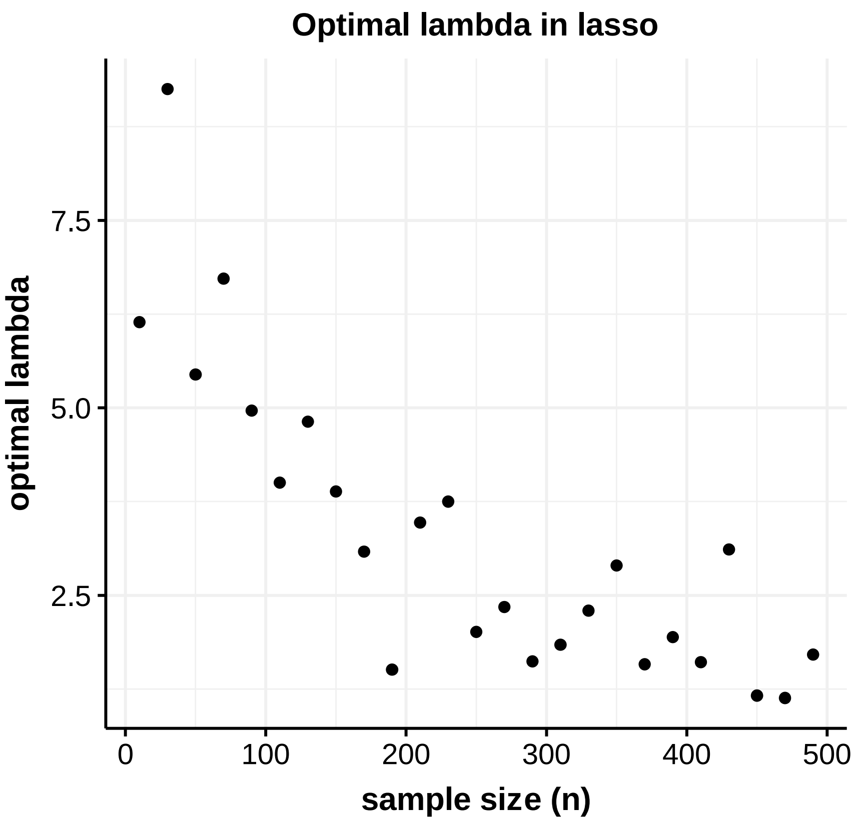

**Supplementary Figure 2: Performance of alternative methods on simulated data generated with different sample sizes and different genetic architectures.** Data are simulated for continuous phenotype under a mild negative selection model and three different degrees of polygenicity (top panel: $p_{causal} = 0.01$, middle panel: $p_{causal} = 0.001$, and bottom panel: $p_{causal} = 5 \times 10^{-4}$). Common SNP heritability is fixed at 0.4 across all populations, and the correlations in effect sizes for share SNPs between all pairs of populations is fixed at 0.8. The sample sizes for GWAS training data are assumed to be (**a**) n=15,000, and (**b**) n=80,000 for the four non-EUR target populations; and is fixed at n=100,000 for the EUR population. PRS generated from all methods are tuned in n=10,000 samples, and then tested in n=10,000 independent samples in each target population. The PRS-CSx package is restricted to SNPs from HM3, whereas other alternative methods use SNPs from either HM3 or MEGA. Bars in the figure show the performance of $R^2$ for each method in each dataset. Colors are described on the right side of the figure. Source data are provided in Supplementary Data 2.

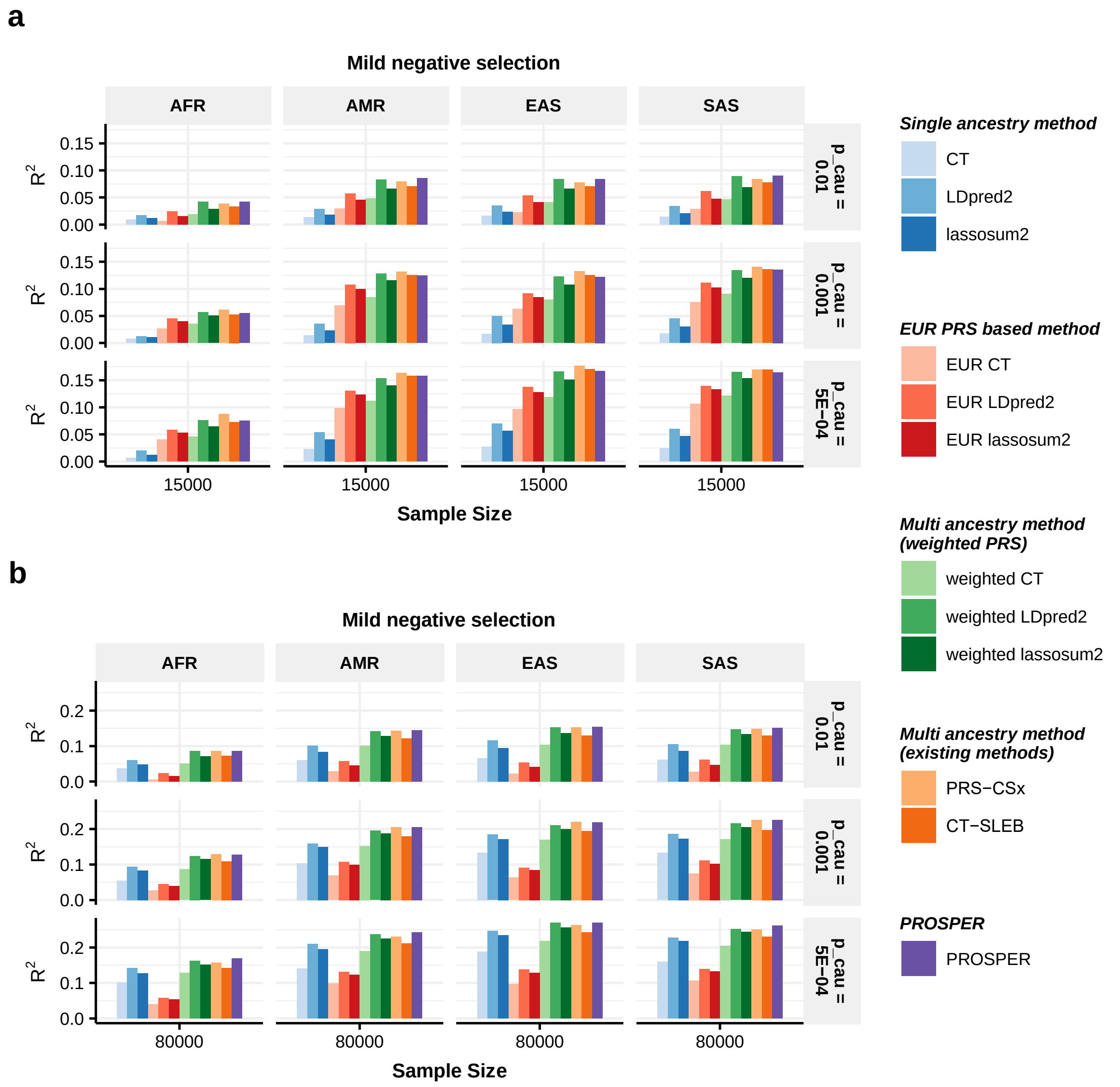

**Supplementary Figure 3: Performance of alternative methods on simulated data generated with different sample sizes and different genetic architectures.** Data are simulated for continuous phenotype under a no negative selection model and three different degrees of polygenicity (top panel: $p_{causal} = 0.01$, middle panel: $p_{causal} = 0.001$, and bottom panel: $p_{causal} = 5 \times 10^{-4}$). Common SNP heritability is fixed at 0.4 across all populations, and the correlations in effect sizes for share SNPs between all pairs of populations is fixed at 0.8. The sample sizes for GWAS training data are assumed to be (**a**) n=15,000, and (**b**) n=80,000 for the four non-EUR target populations; and is fixed at n=100,000 for the EUR population. PRS generated from all methods are tuned in n=10,000 samples, and then tested in n=10,000 independent samples in each target population. The PRS-CSx package is restricted to SNPs from HM3, whereas other alternative methods use SNPs from either HM3 or MEGA. Bars in the figure show the performance of $R^2$ for each method in each dataset. Colors are described on the right side of the figure. Source data are provided in Supplementary Data 3.

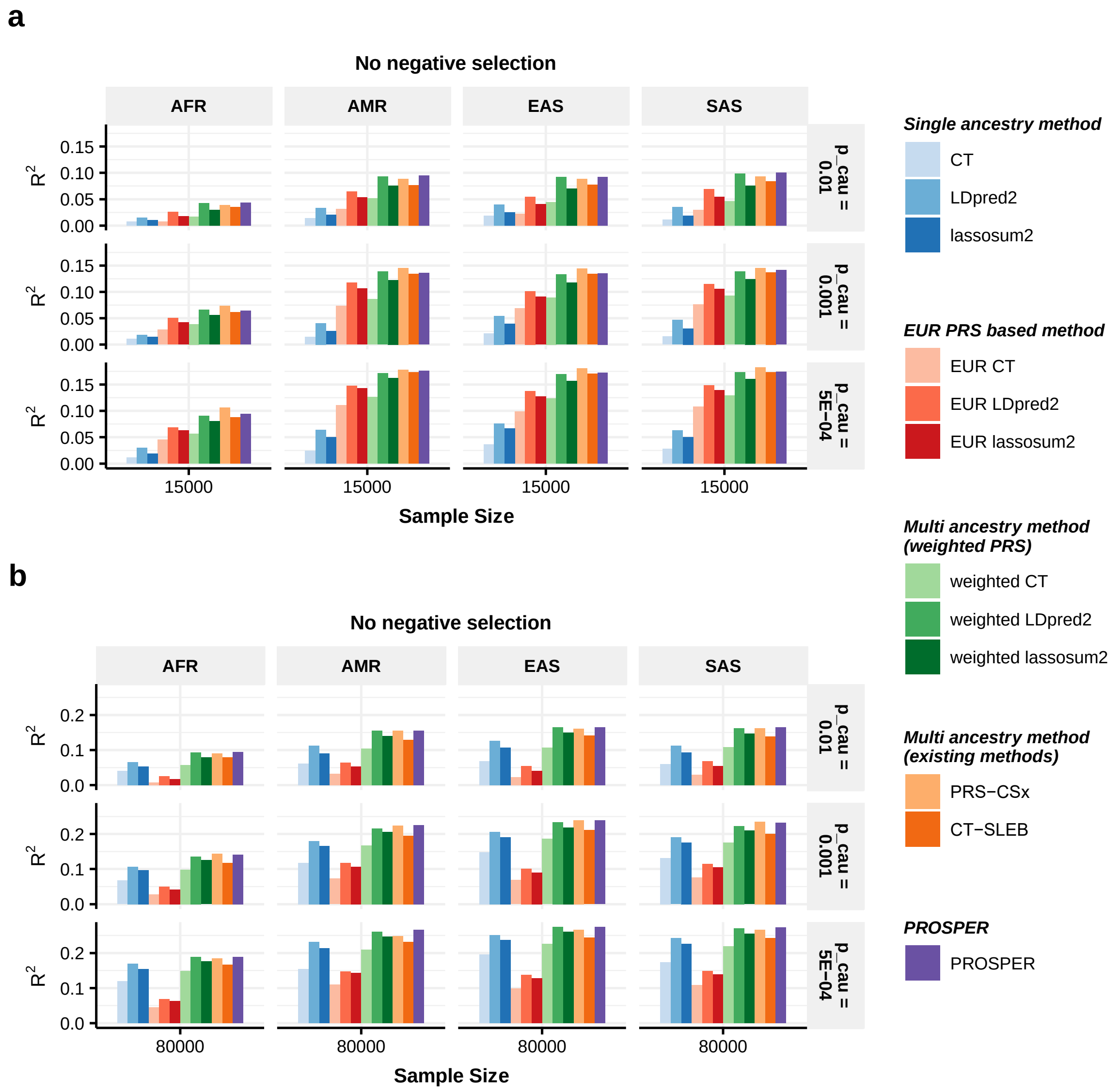

**Supplementary Figure 4: Performance of alternative methods on simulated data generated with different sample sizes and different genetic architectures.** Data are simulated for continuous phenotype under a strong negative selection model and three different degrees of polygenicity (top panel: $p_{causal} = 0.01$, middle panel: $p_{causal} = 0.001$, and bottom panel: $p_{causal} = 5 \times 10^{-4}$). Per-SNP heritability is assumed to be the same across all populations and thus leads to the common SNP heritability value of 0.32, 0.21, 0.16, 0.19 and 0.17 for AFR, AMR, EAS, EUR and SAS, respectively. The correlations in effect sizes for share SNPs between all pairs of populations is fixed at 0.8. The sample sizes for GWAS training data are assumed to be (**a**) n=15,000, and (**b**) n=80,000 for the four non-EUR target populations; and is fixed at n=100,000 for the EUR population. PRS generated from all methods are tuned in n=10,000 samples, and then tested in n=10,000 independent samples in each target population. The PRS-CSx package is restricted to SNPs from HM3, whereas other alternative methods use SNPs from either HM3 or MEGA. Bars in the figure show the performance of $R^2$ for each method in each dataset. Colors are described on the right side of the figure. Source data are provided in Supplementary Data 4.

**a**

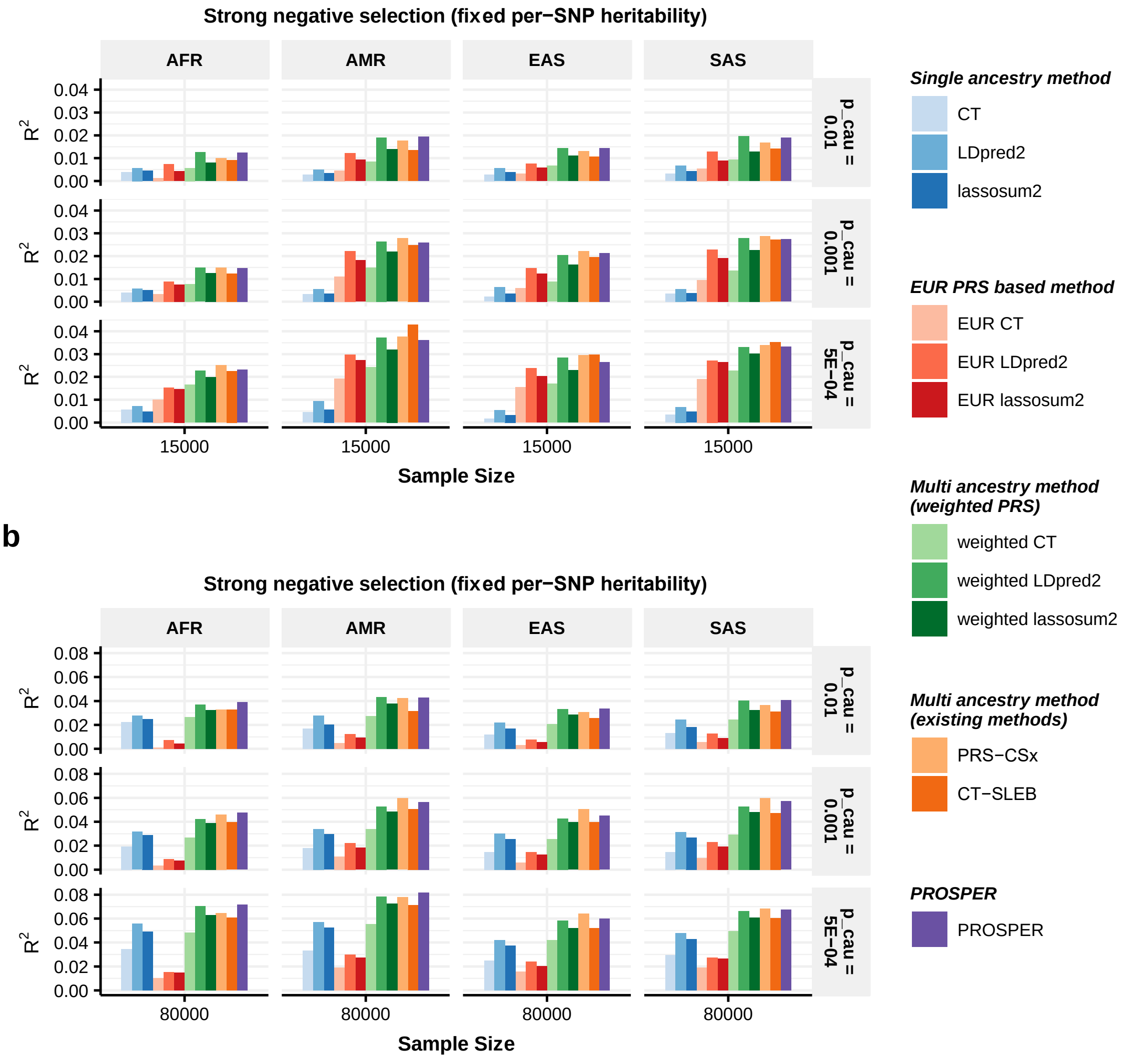


**b**

**Supplementary Figure 5: Performance of alternative methods on simulated data generated with different sample sizes and different genetic architectures.** Data are simulated for continuous phenotype under a <u>strong negative selection</u> model and three different degrees of polygenicity (top panel: $p_{causal} = 0.01$, middle panel: $p_{causal} = 0.001$, and bottom panel: $p_{causal} = 5 \times 10^{-4}$). <u>Per-SNP heritability is assumed to be the same across all populations, and the correlations in effect sizes for share SNPs between all pairs of populations is fixed at 0.6</u>. The sample sizes for GWAS training data are assumed to be (**a**) n=15,000, and (**b**) n=80,000 for the four non-EUR target populations; and is fixed at n=100,000 for the EUR population. PRS generated from all methods are tuned in n=10,000 samples, and then tested in n=10,000 independent samples in each target population. The PRS-CSx package is restricted to SNPs from HM3, whereas other alternative methods use SNPs from either HM3 or MEGA. Bars in the figure show the performance of $R^2$ for each method in each dataset. Colors are described on the right side of the figure. Source data are provided in Supplementary Data 5.

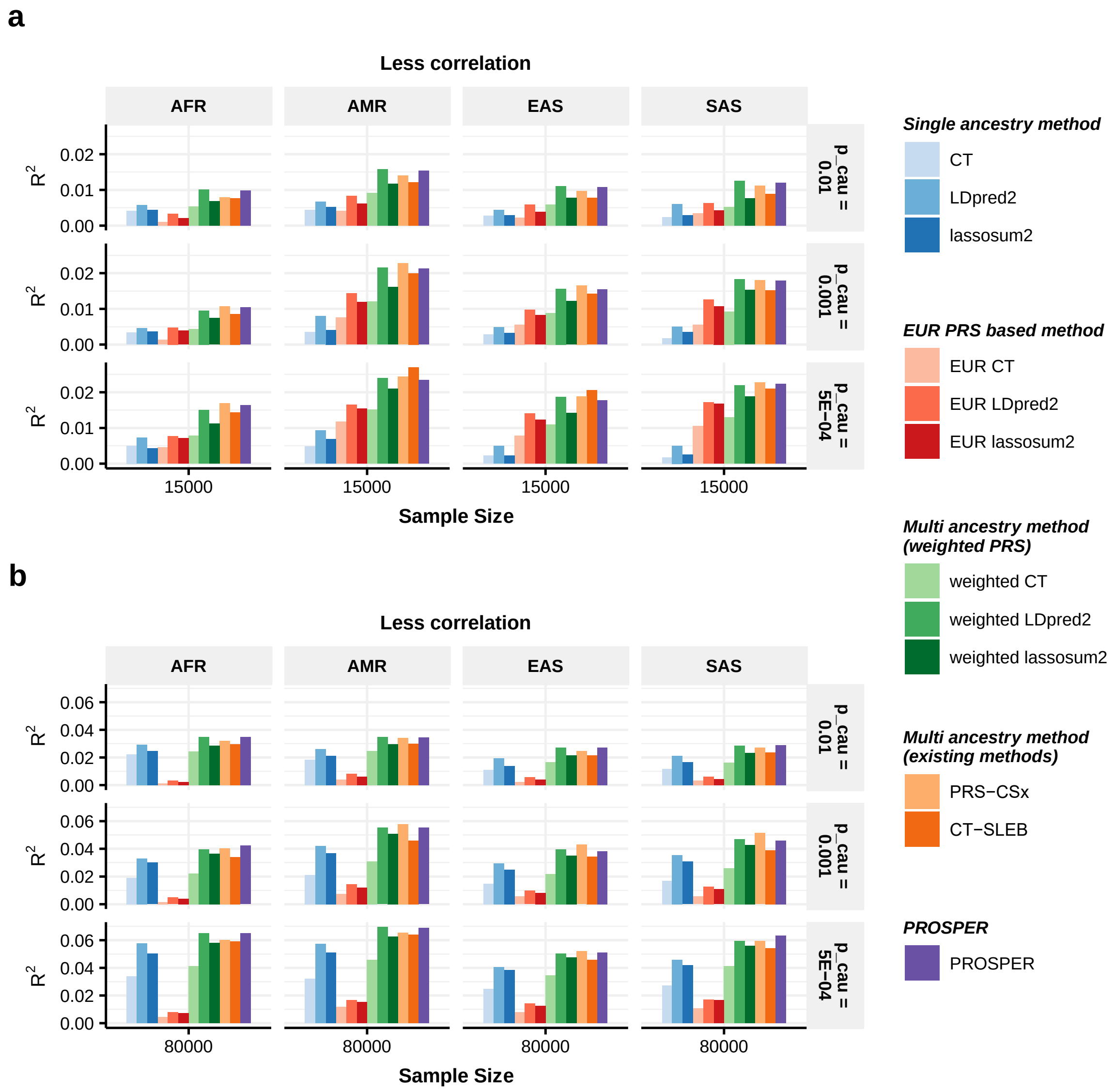

**Supplementary Figure 6: Performance comparison of advanced weighted lassosum2 (with super learning across all tuning parameters and ancestries) and PROSPER for prediction of simulated data generated with different sample sizes and genetic architectures under strong negative selection and fixed common-SNP heritability.** Data used to generate this figure is same as in Figure 2. Bars in the figure show the performance of $R^2$ for each method in each dataset. Colors are described on the right side of the figure. Source data are provided in Supplementary Data 15.

**a**

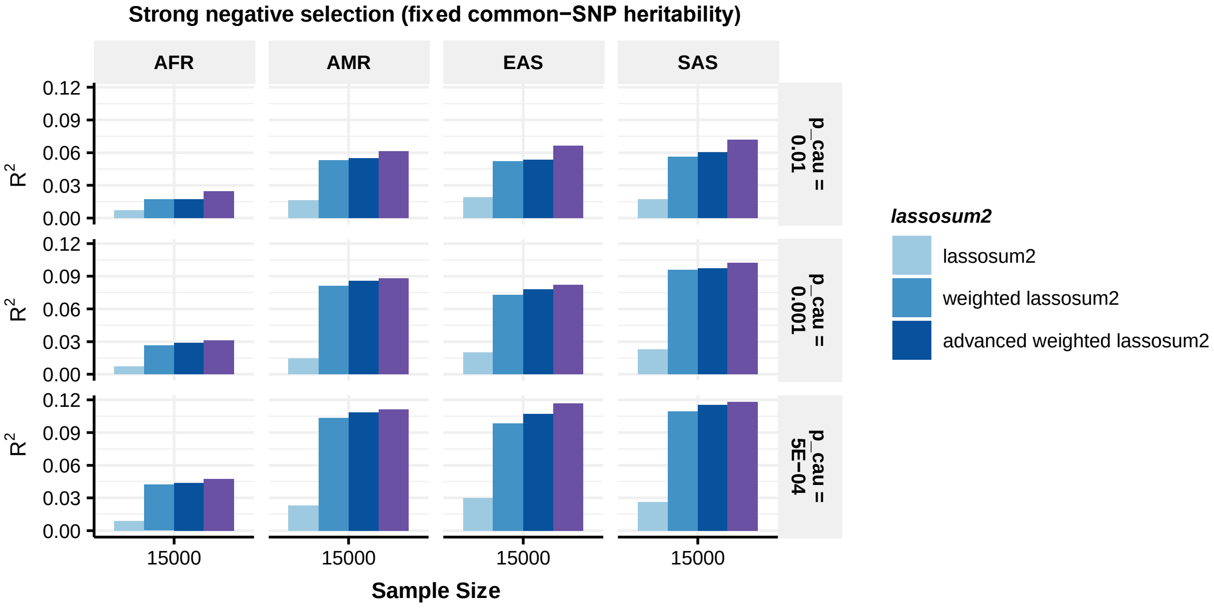


**b**

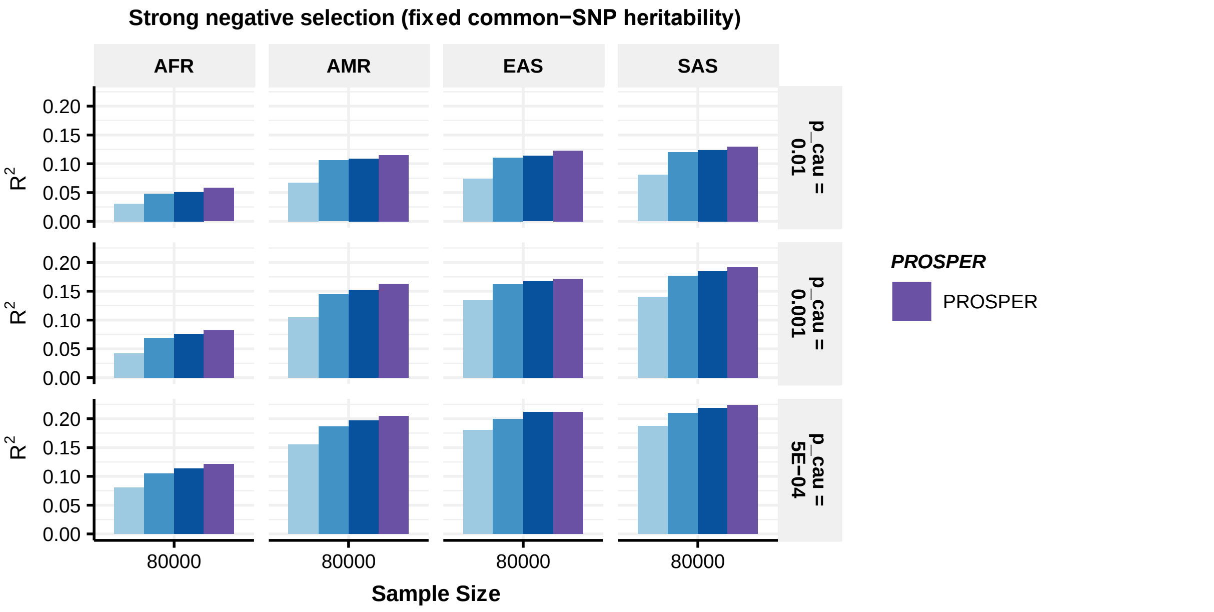

**Supplementary Figure 7: Performance comparison of advanced weighted lassosum2 (with super learning across all tuning parameters and ancestries) and PROSPER for prediction of simulated data generated with different sample sizes and different genetic architectures under mild negative selection.** Data used to generate this figure is same as in Supplementary Figure 2. Bars in the figure show the performance of $R^2$ for each method in each dataset. Colors are described on the right side of the figure. Source data are provided in Supplementary Data 15.

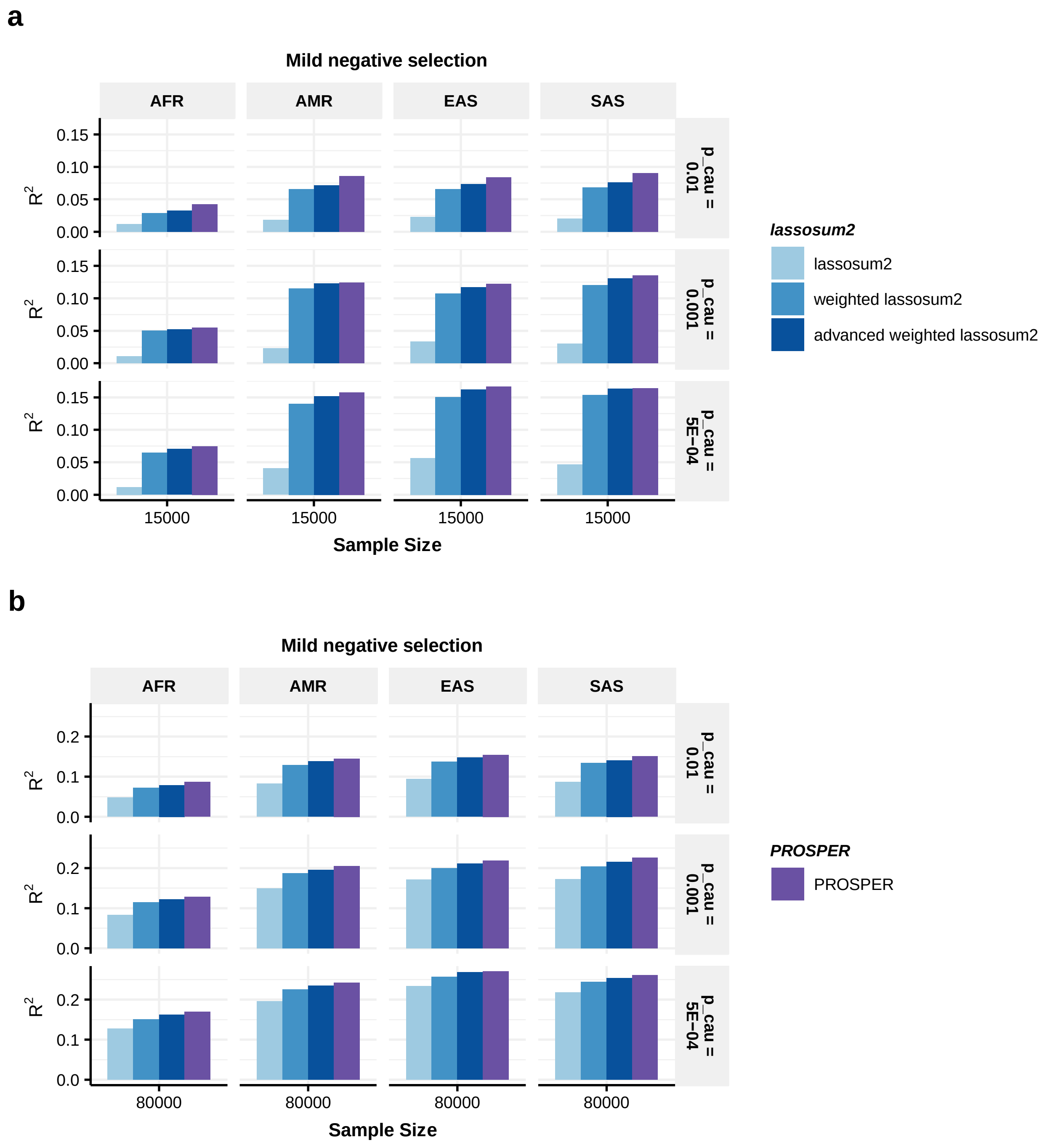

**Supplementary Figure 8: Performance comparison of advanced weighted lassosum2 (with super learning across all tuning parameters and ancestries) and PROSPER for prediction of simulated data generated with different sample sizes and different genetic architectures under no negative selection.** Data used to generate this figure is same as in Supplementary Figure 3. Bars in the figure show the performance of $R^2$ for each method in each dataset. Colors are described on the right side of the figure. Source data are provided in Supplementary Data 15.

**a**

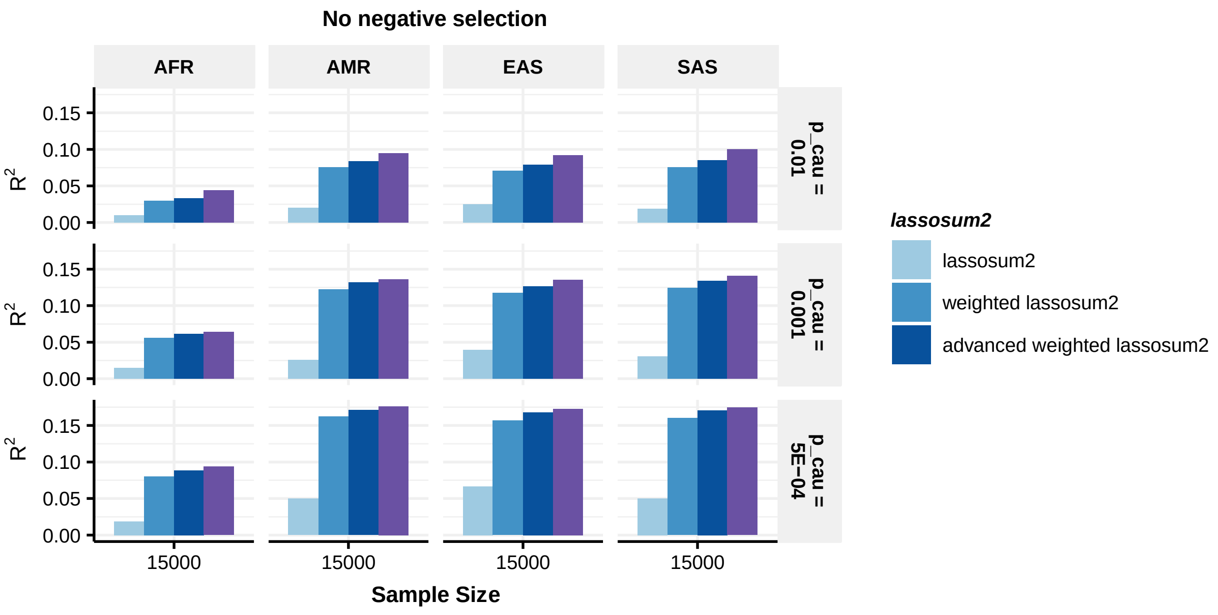


**b**

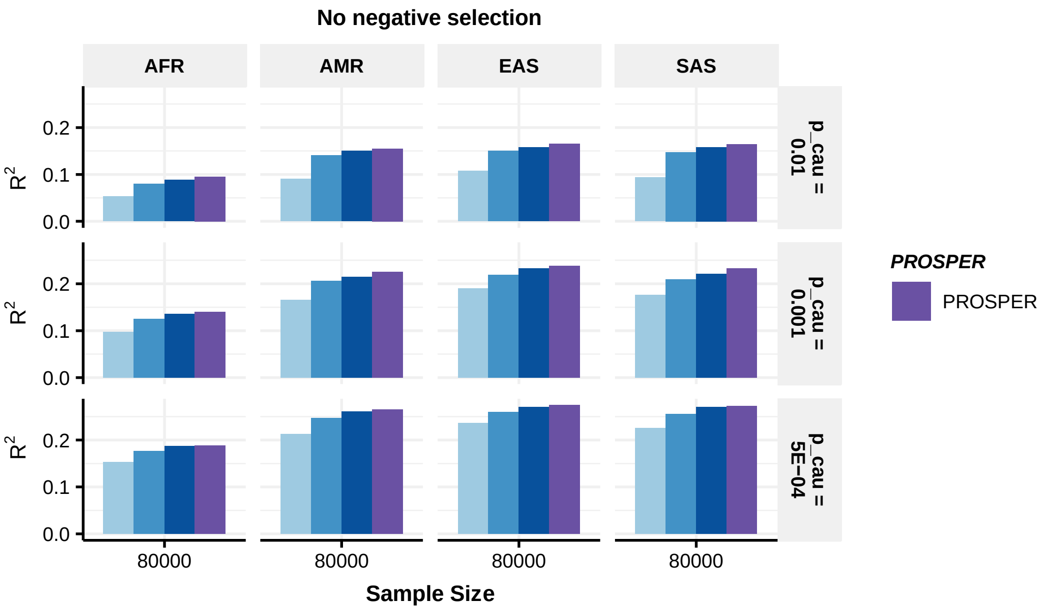

**Supplementary Figure 9: Performance comparison of advanced weighted lassosum2 (with super learning across all tuning parameters and ancestries) and PROSPER for prediction of simulated data generated with different sample sizes and genetic architectures under strong negative selection and fixed per-SNP heritability.** Data used to generate this figure is same as in Supplementary Figure 4. Bars in the figure show the performance of $R^2$ for each method in each dataset. Colors are described on the right side of the figure. Source data are provided in Supplementary Data 15.

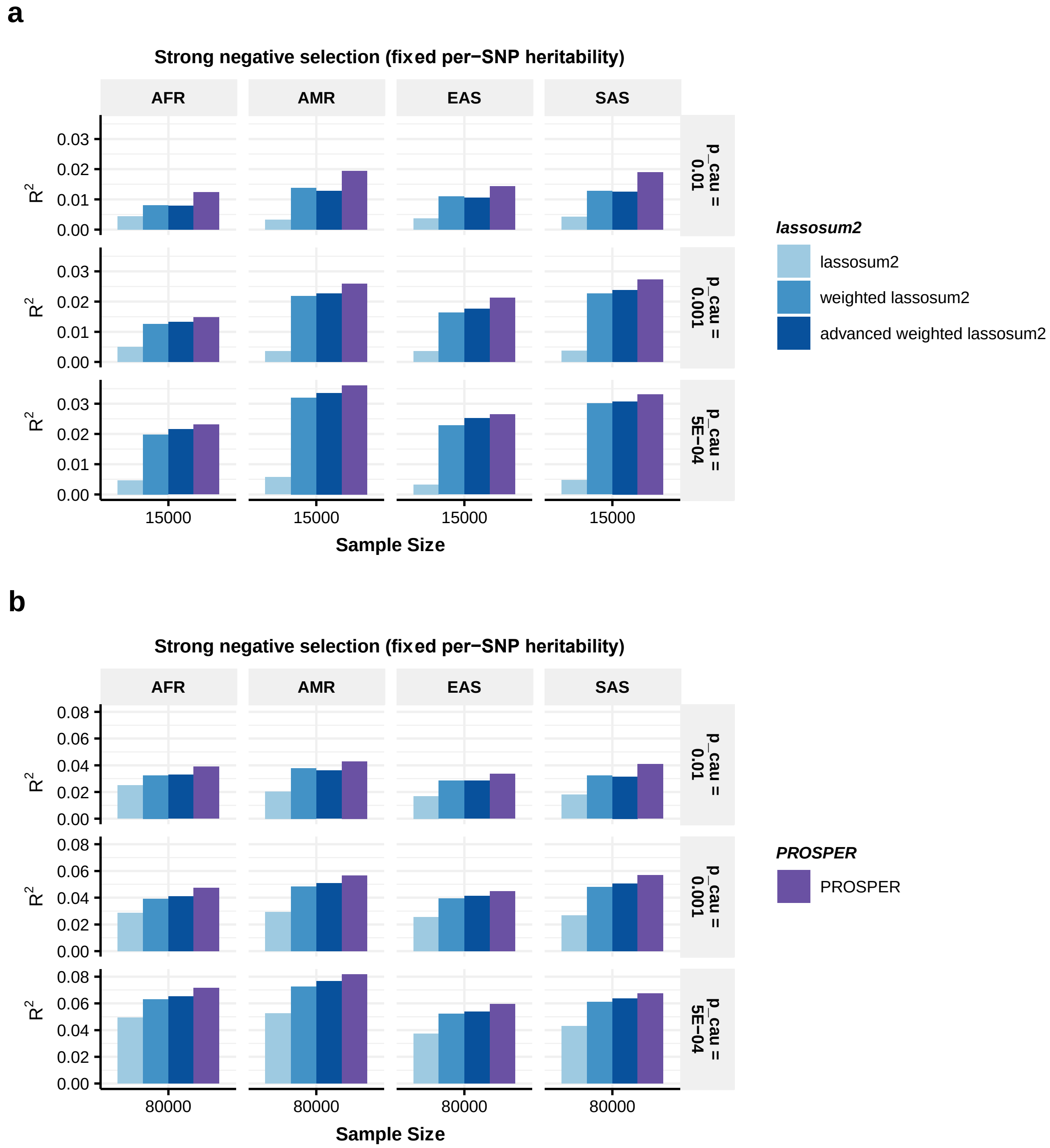

**Supplementary Figure 10: Performance comparison of advanced weighted lassosum2 (with super learning across all tuning parameters and ancestries) and PROSPER for prediction of simulated data generated with different sample sizes and genetic architectures under strong negative selection and less genetic correlation.** Data used to generate this figure is same as in Supplementary Figure 5. Bars in the figure show the performance of $R^2$ for each method in each dataset. Colors are described on the right side of the figure. Source data are provided in Supplementary Data 15.

**a**

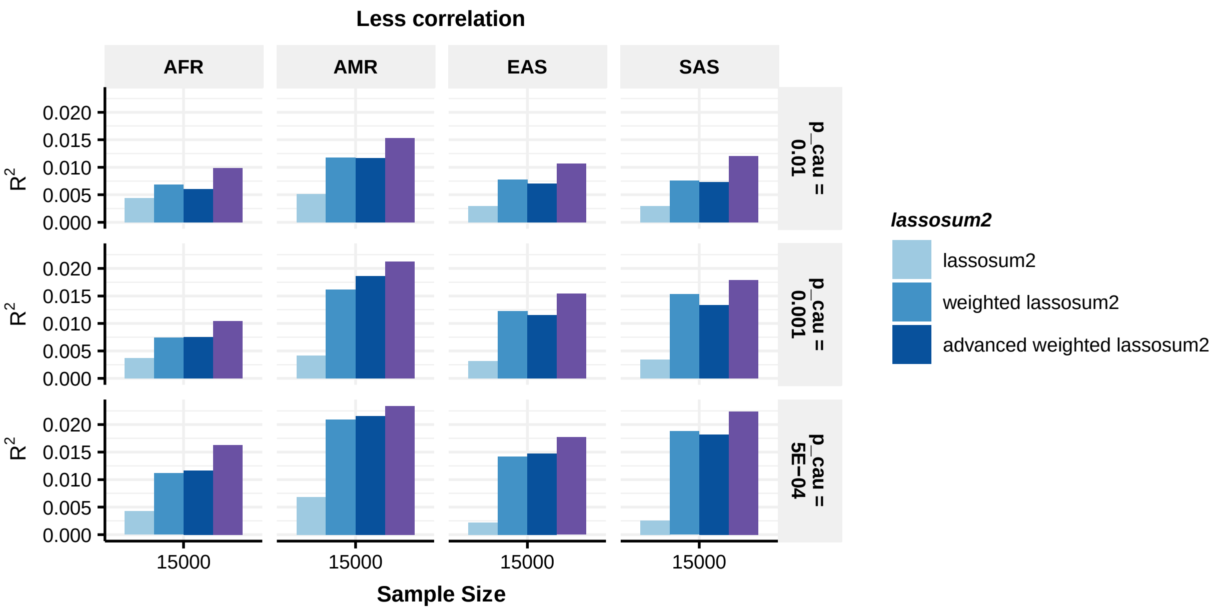


**b**

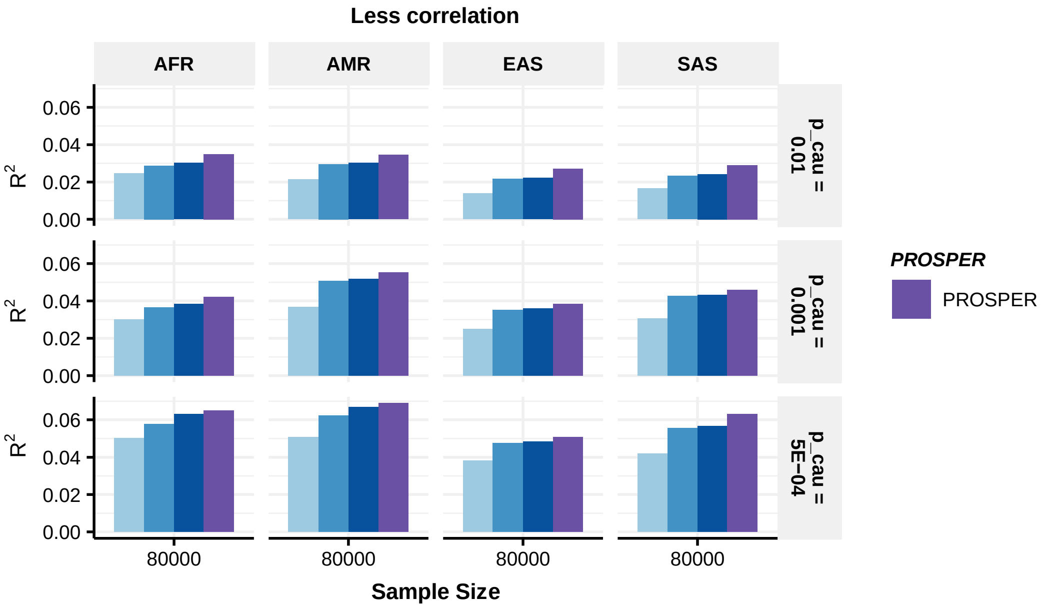

**Supplementary Figure 11: Performance comparison of advanced weighted lassosum2 (with super learning across all tuning parameters and ancestries) and PROSPER for prediction of four blood lipid traits (GLGC-training and UKBB-tuning/validation)**. Data used to generate this figure is same as in Figure 5. Bars in the figure show the performance of adjusted $R^2$ for each method in each dataset. Colors are described on the right side of the figure. Source data are provided in Supplementary Data 16.

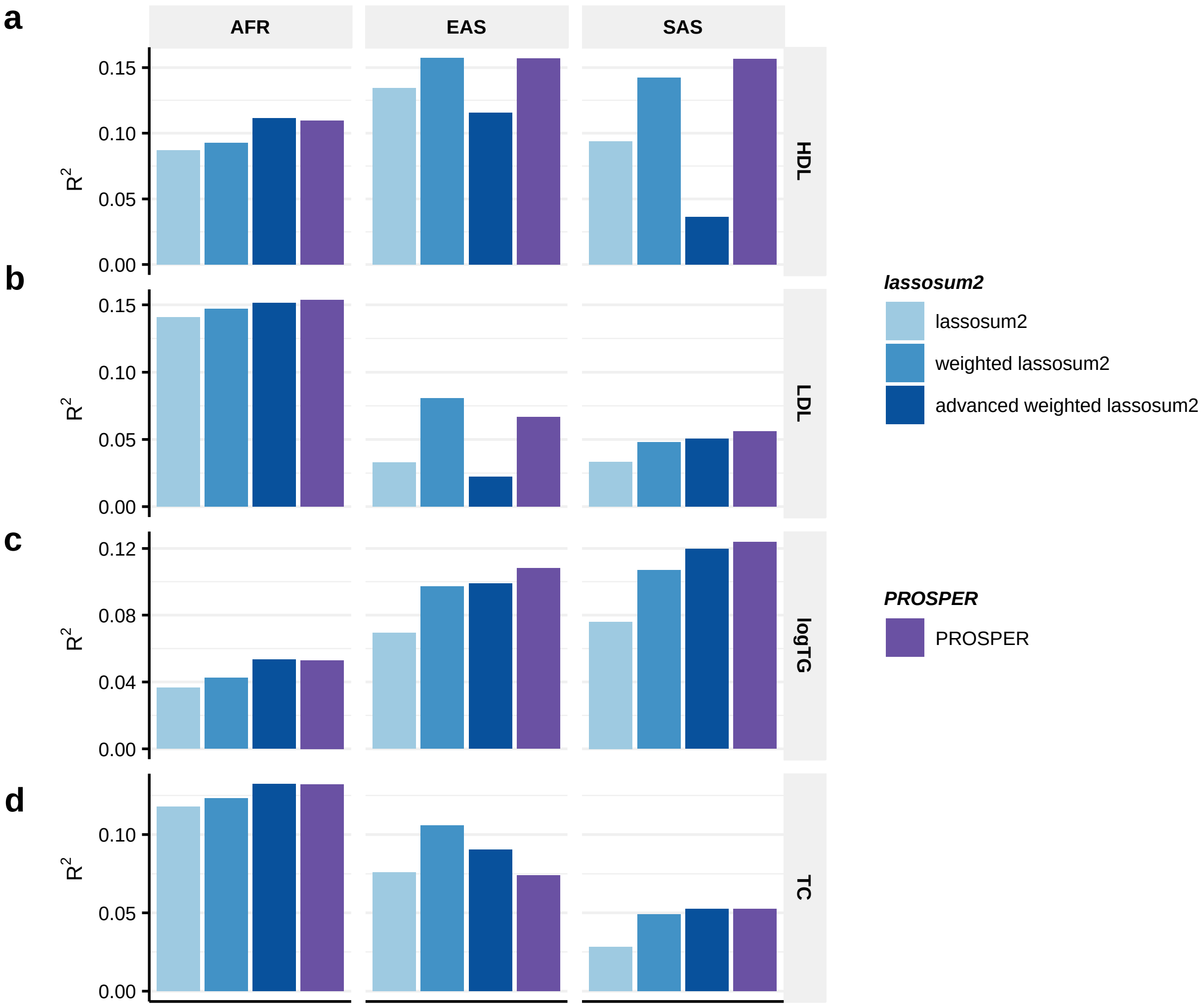

**Supplementary Figure 12: Performance comparison of advanced weighted lassosum2 (with super learning across all tuning parameters and ancestries) and PROSPER for prediction of of two anthropometric traits (AoU-training and UKBB-tuning/validation)**. Data used to generate this figure is same as in Figure 6. Bars in the figure show the performance of adjusted $R^2$ for each method in each dataset. Colors are described on the right side of the figure. Source data are provided in Supplementary Data 16.

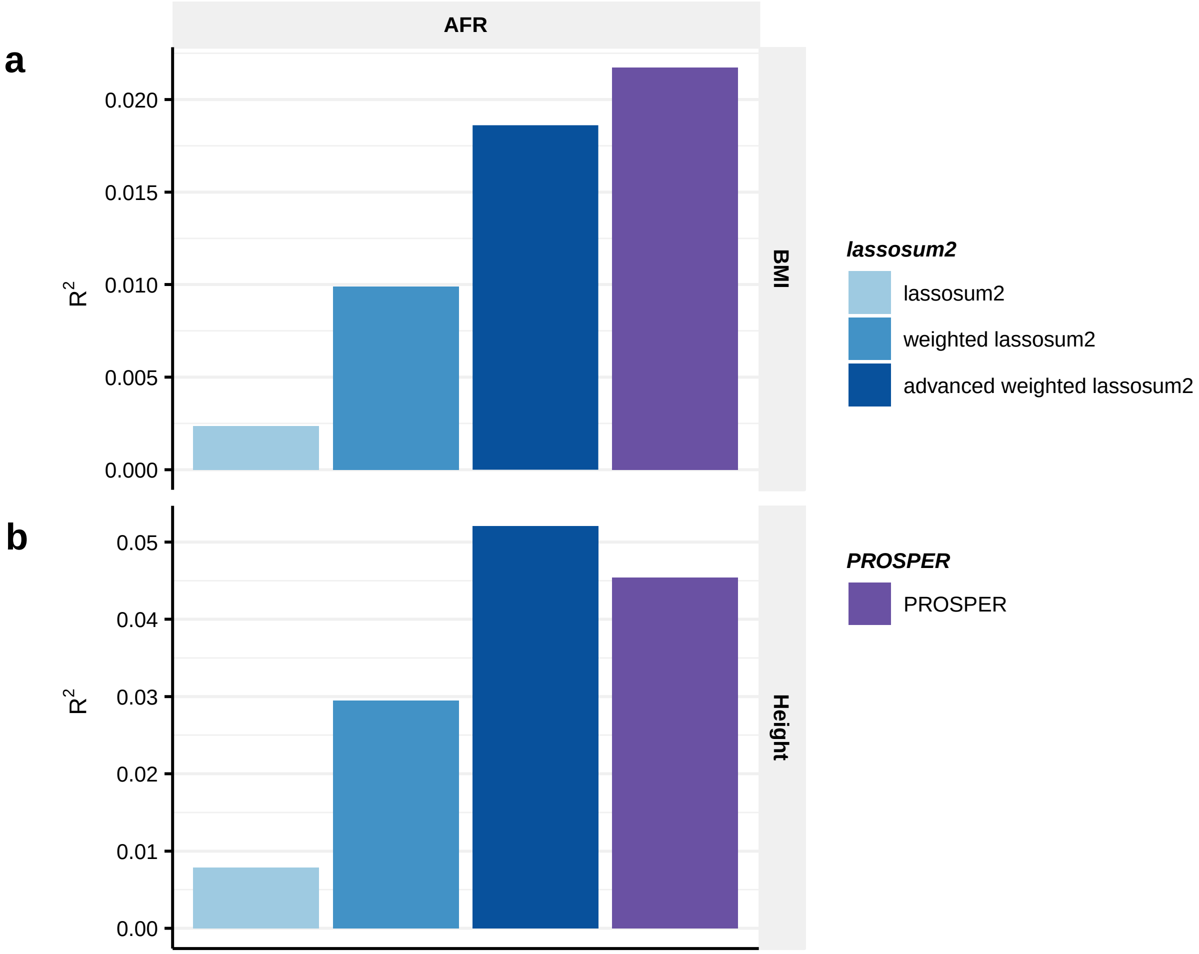

**Supplementary Figure 13: The relationship between tuning sample size and predictive $R^2$.** Data used to generate this figure is same as in Figure 2. PRS is tuned with n=5000, n=3000, n=1000, n=500, n=300, and n=100 tuning samples, and then tested in n=10,000 independent samples in each target population. Colors are described on the right side of the figure. Source data are provided as a Source Data file.

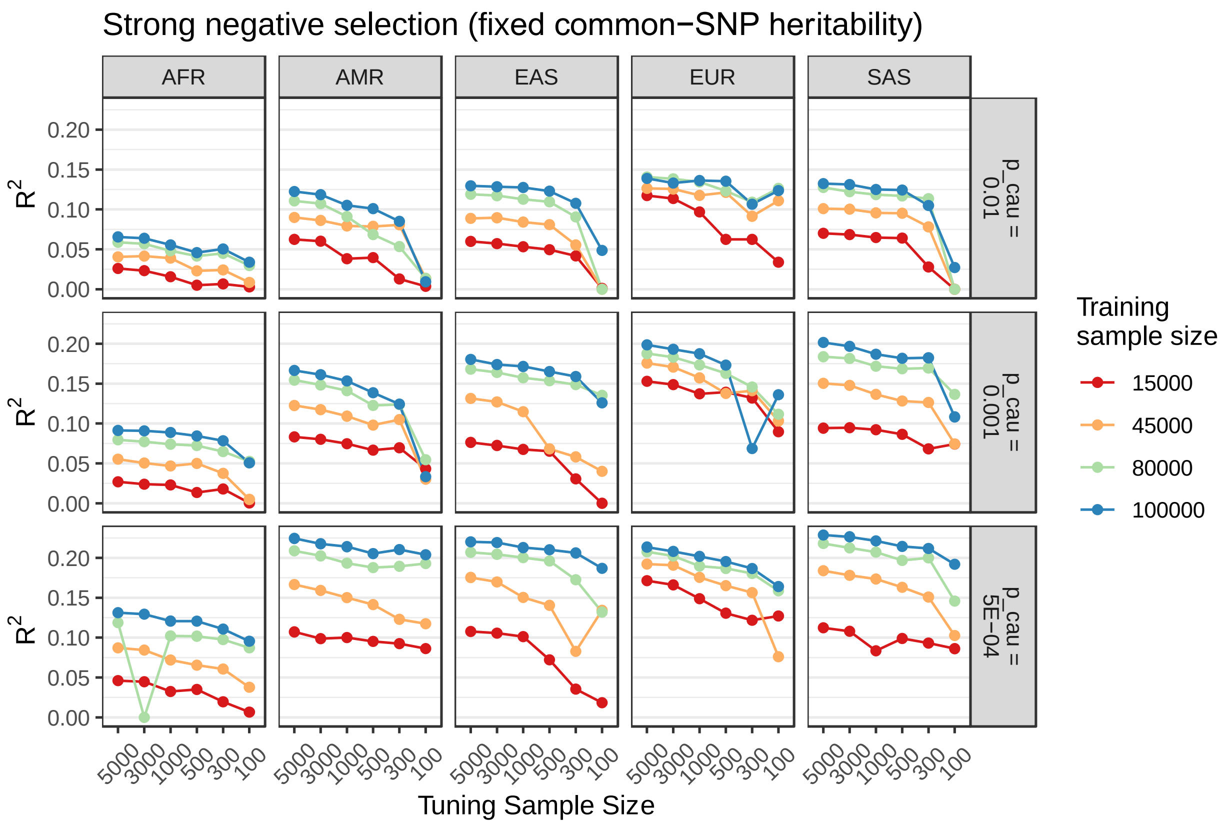

**Full list of members in 23andMe Research Team**

Jianan Zhan, Jared O’Connell, Yunxuan Jiang, Stella Aslibekyan, Adam Auton, Elizabeth Babalola, Robert K. Bell, Jessica Bielenberg, Katarzyna Bryc, Emily Bullis, Daniella Coker, Gabriel Cuellar Partida, Devika Dhamija, Sayantan Das, Sarah L. Elson, Nicholas Eriksson, Teresa Filshtein, Alison Fitch, Kipper Fletez-Brant, Pierre Fontanillas, Will Freyman, Julie M. Granka, Karl Heilbron, Alejandro Hernandez, Barry Hicks, David A. Hinds, Ethan M. Jewett, Katelyn Kukar, Alan Kwong, Keng-Han Lin, Bianca A. Llamas, Maya Lowe, Jey C. McCreight, Matthew H. McIntyre, Steven J. Micheletti, Meghan E. Moreno, Priyanka Nandakumar, Dominique T. Nguyen, Elizabeth S. Noblin, Aaron A. Petrakovitz, G. David Poznik, Alexandra Reynoso, Morgan Schumacher, Anjali J. Shastri, Janie F. Shelton, Jingchunzi Shi, Suyash Shringarpure, Qiaojuan Jane Su, Susana A. Tat, Christophe Toukam Tchakouté, Vinh Tran, Joyce Y. Tung, Xin Wang, Wei Wang, Catherine H. Weldon, Peter Wilton, Corinna D. Wong & Bertram L. Koelsch
